## Supplementary figures and images for "Zyxin contributes to coupling between cell junctions and contractile actomyosin networks during apical constriction"

### S-Fig_1.tif

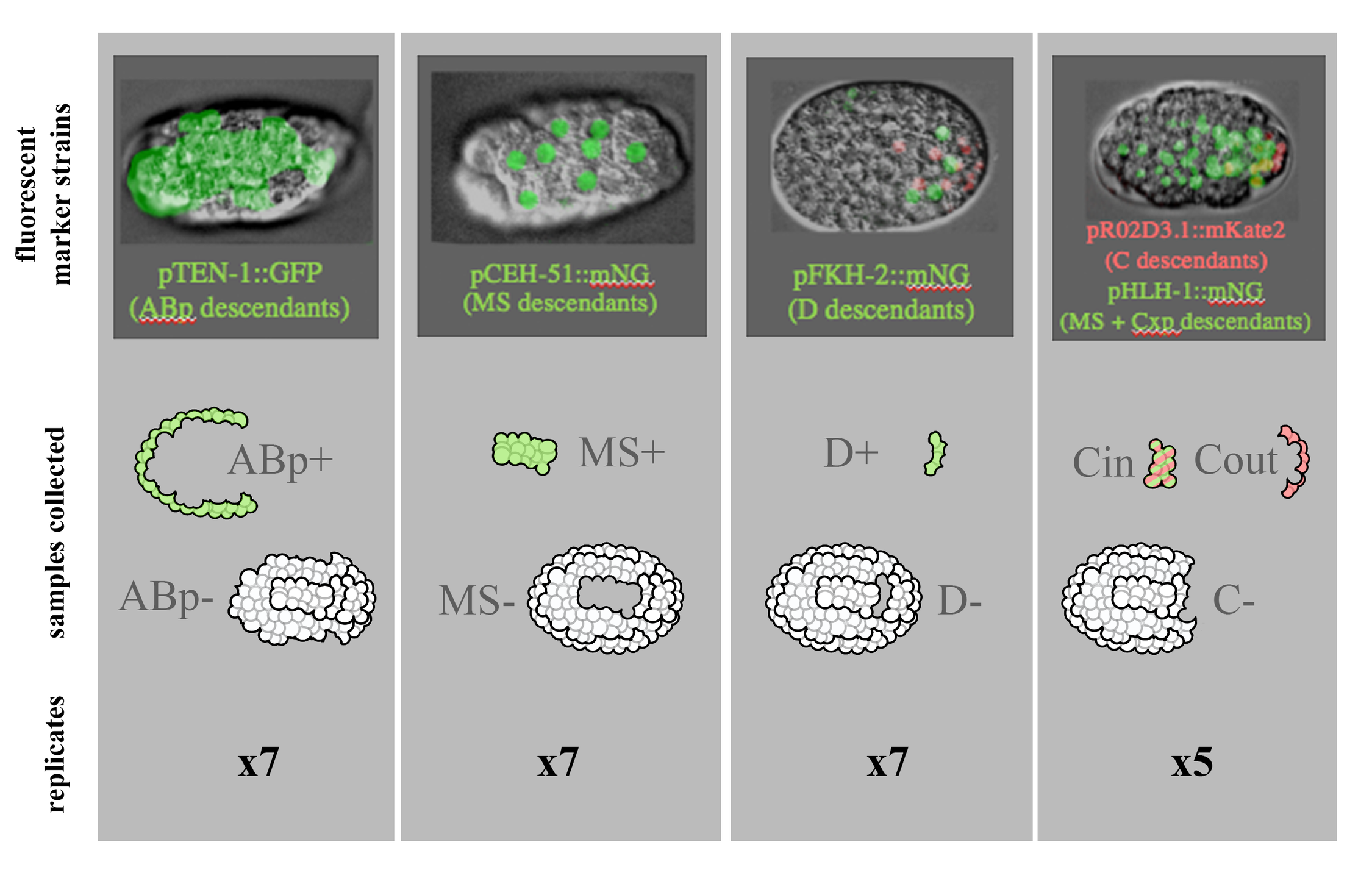

### S-Fig_2.tif

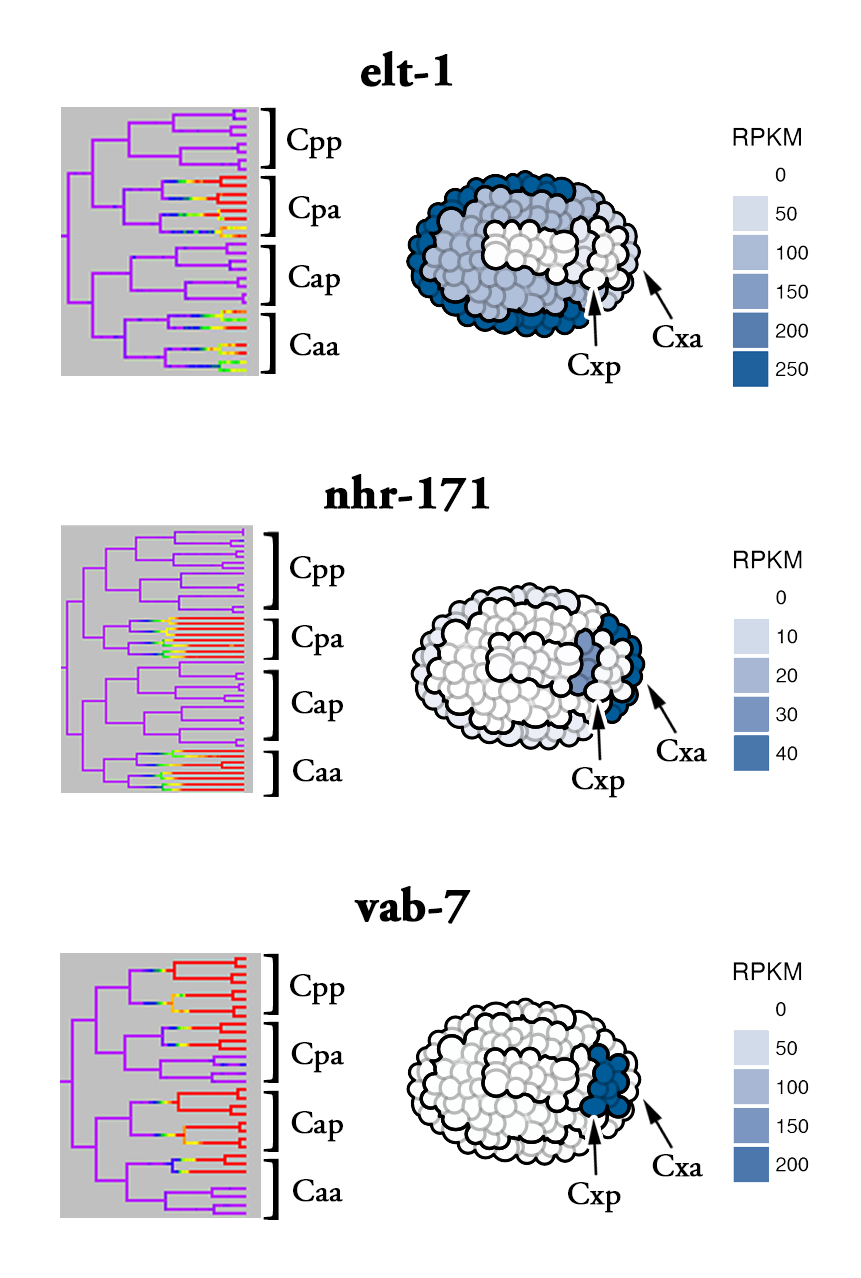

### S-Fig_3.tif

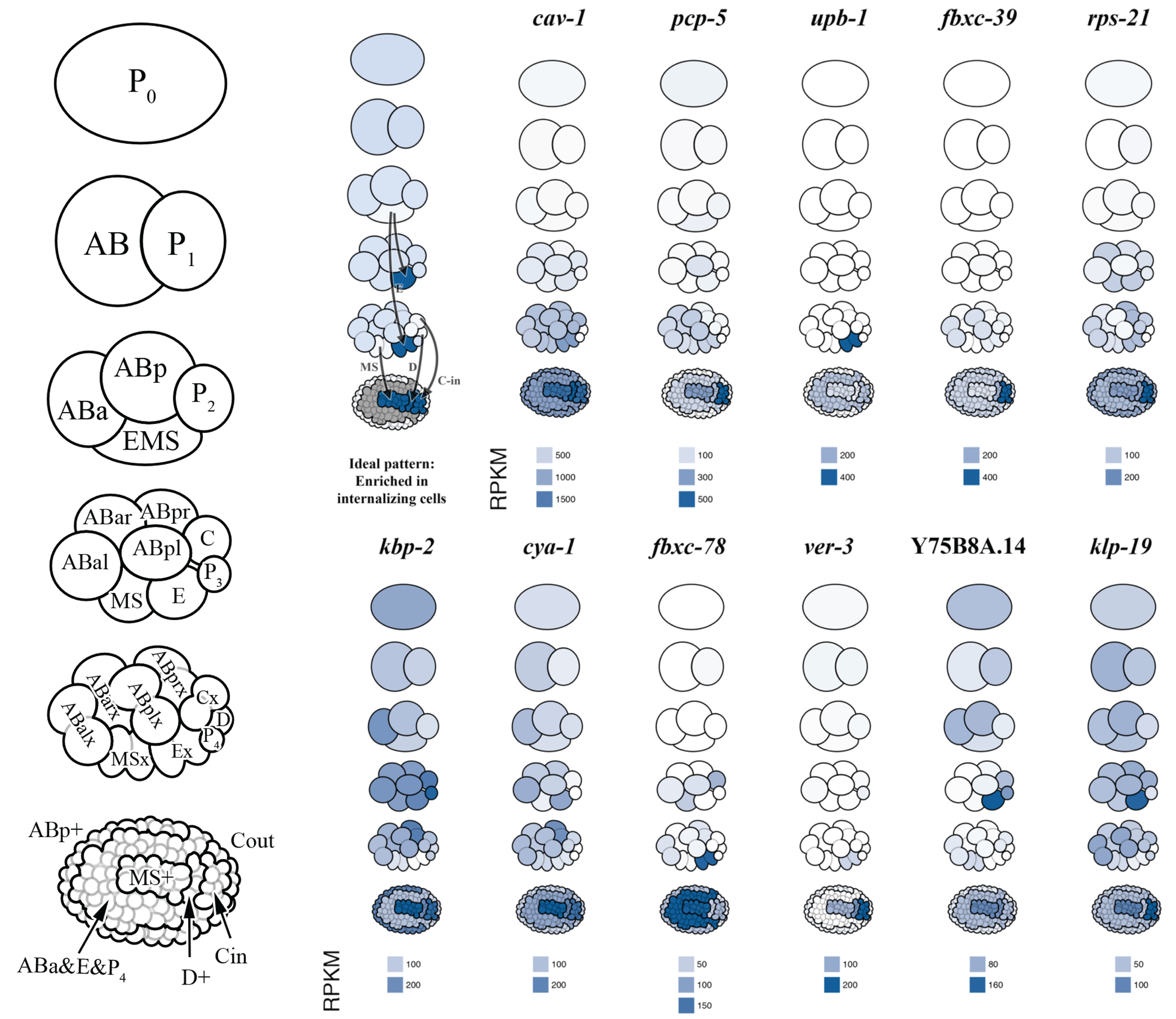

### S-Fig_4.tif

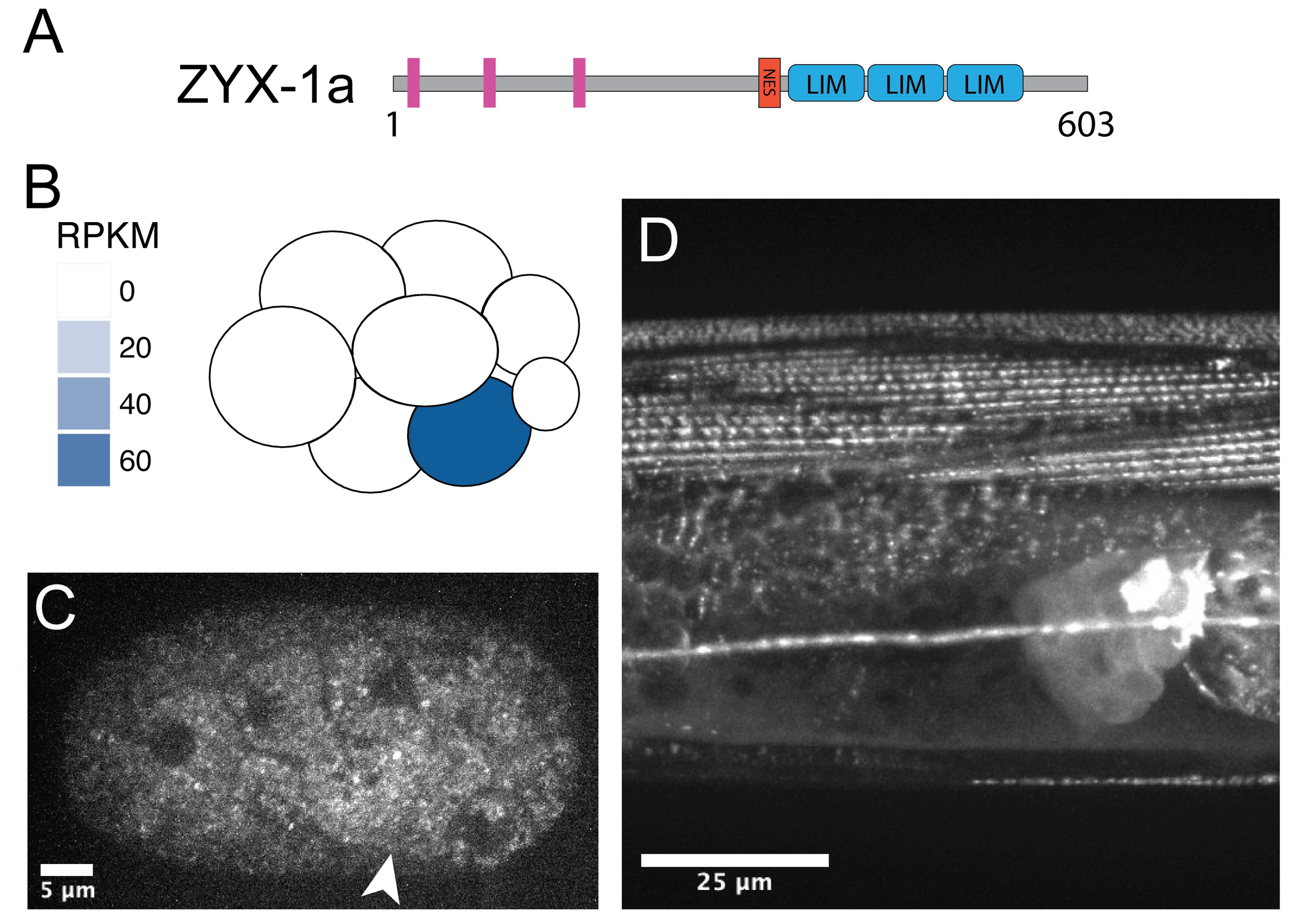

### S-Fig_5.tif

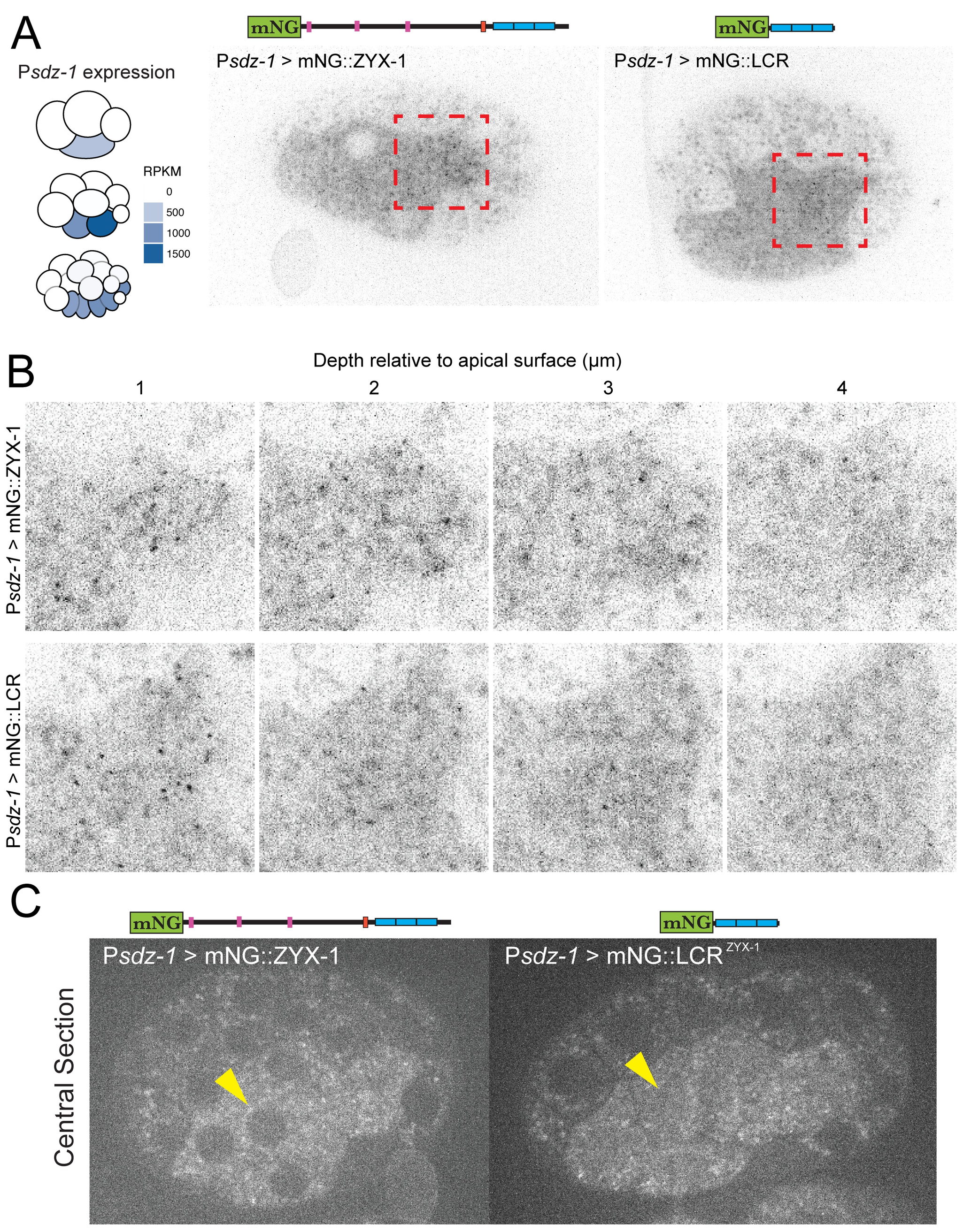

### S-Fig_6.tif

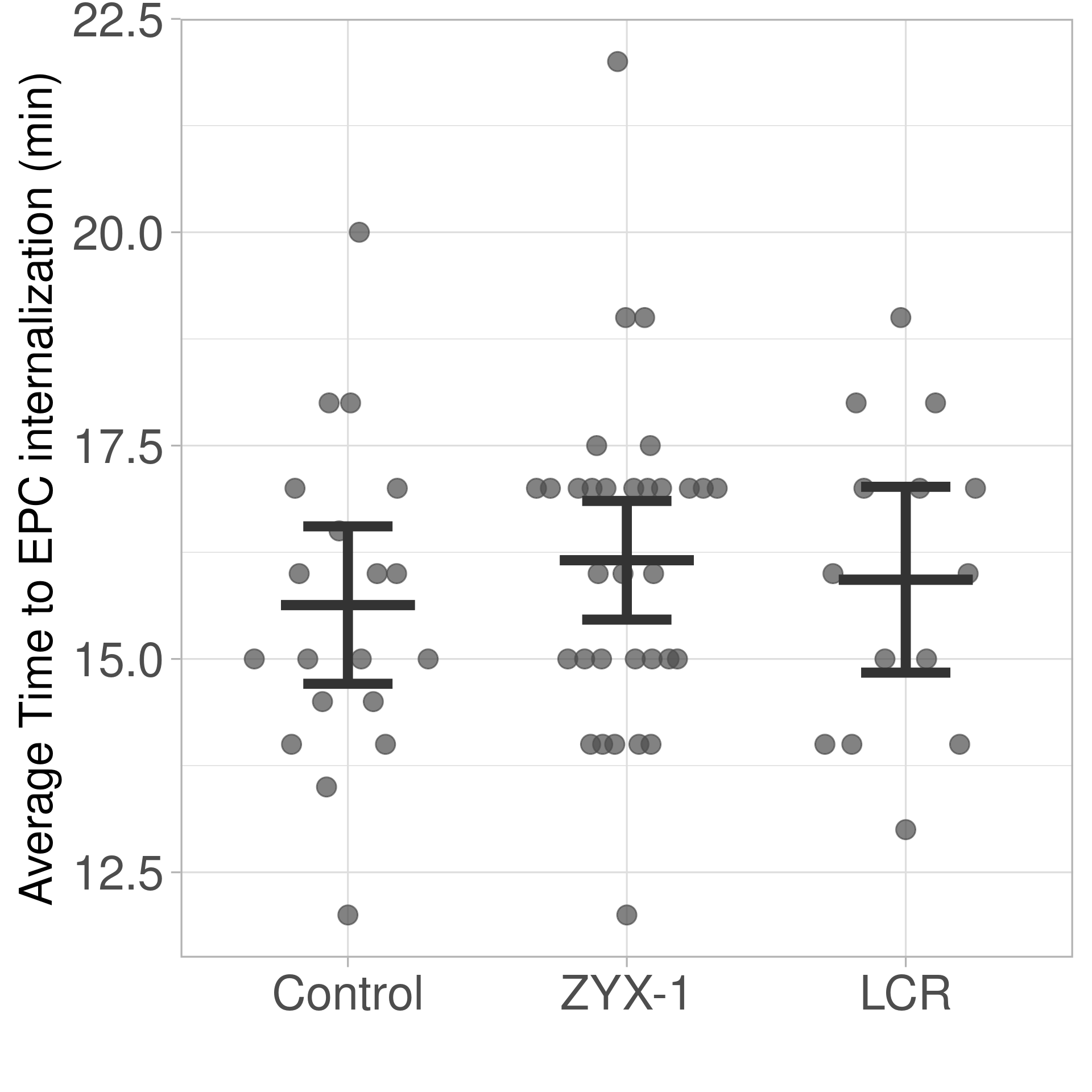

### S-Table_01.tif

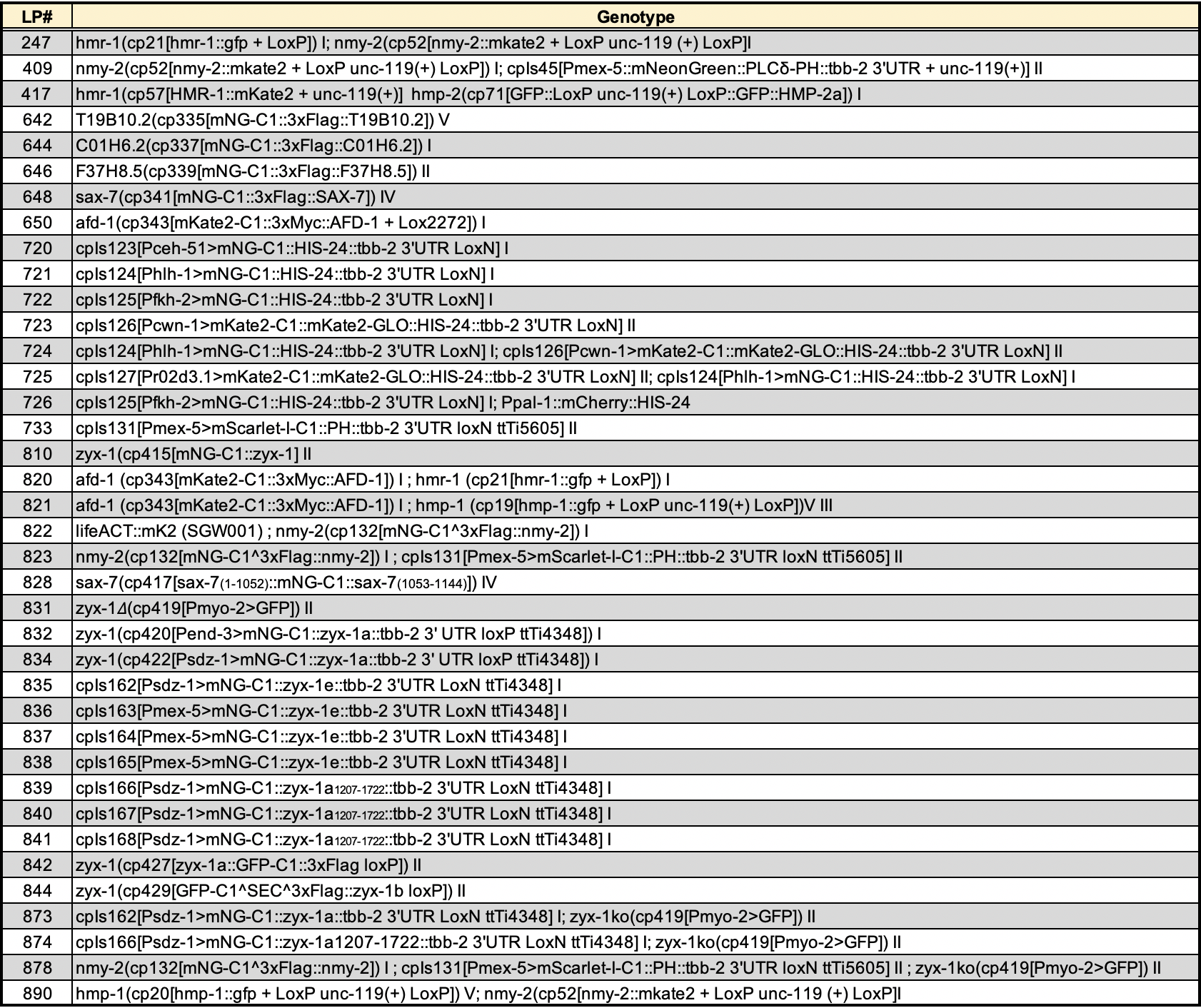

### S-Table_02.tif

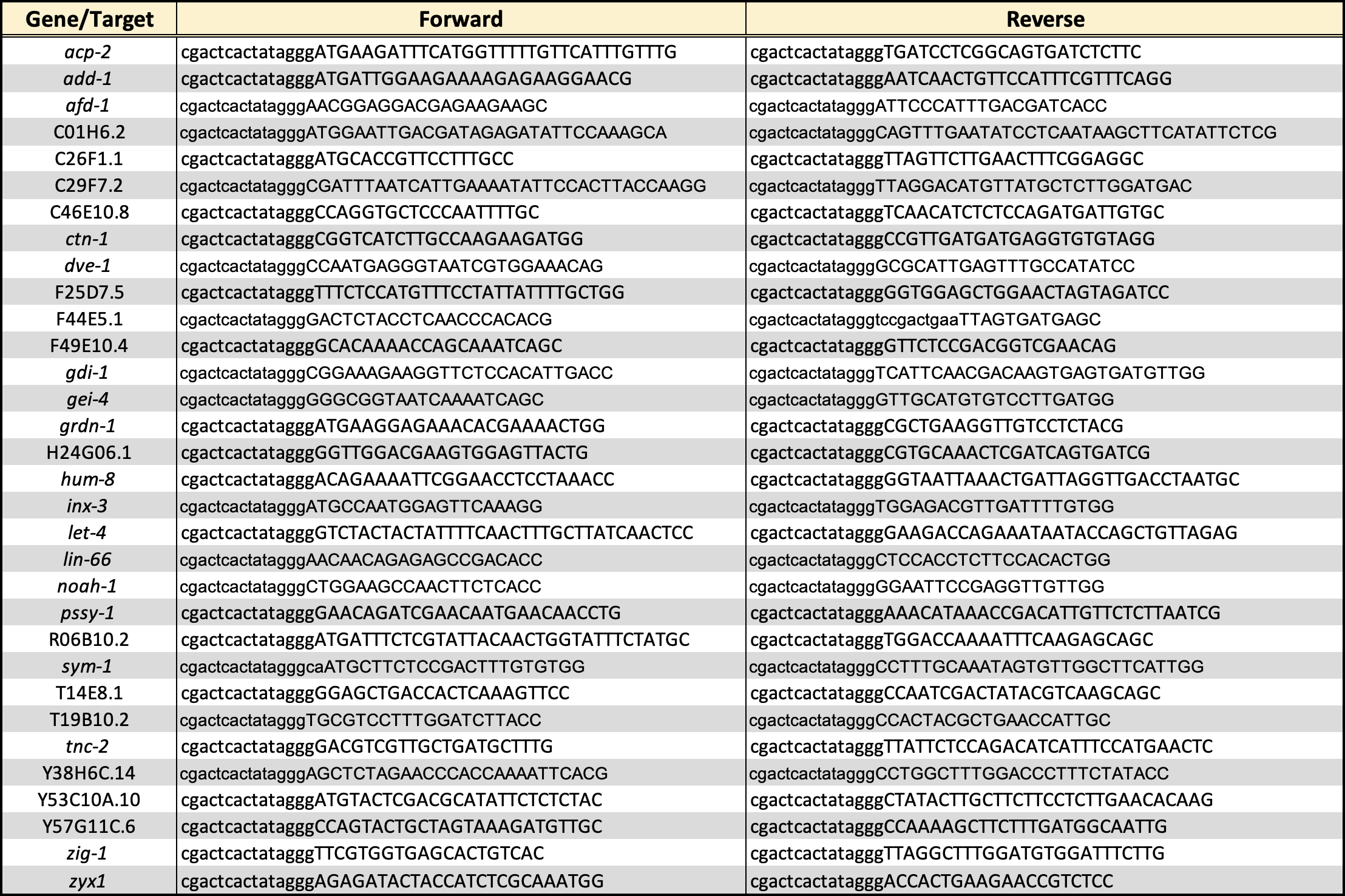

### S-Table_3.tif

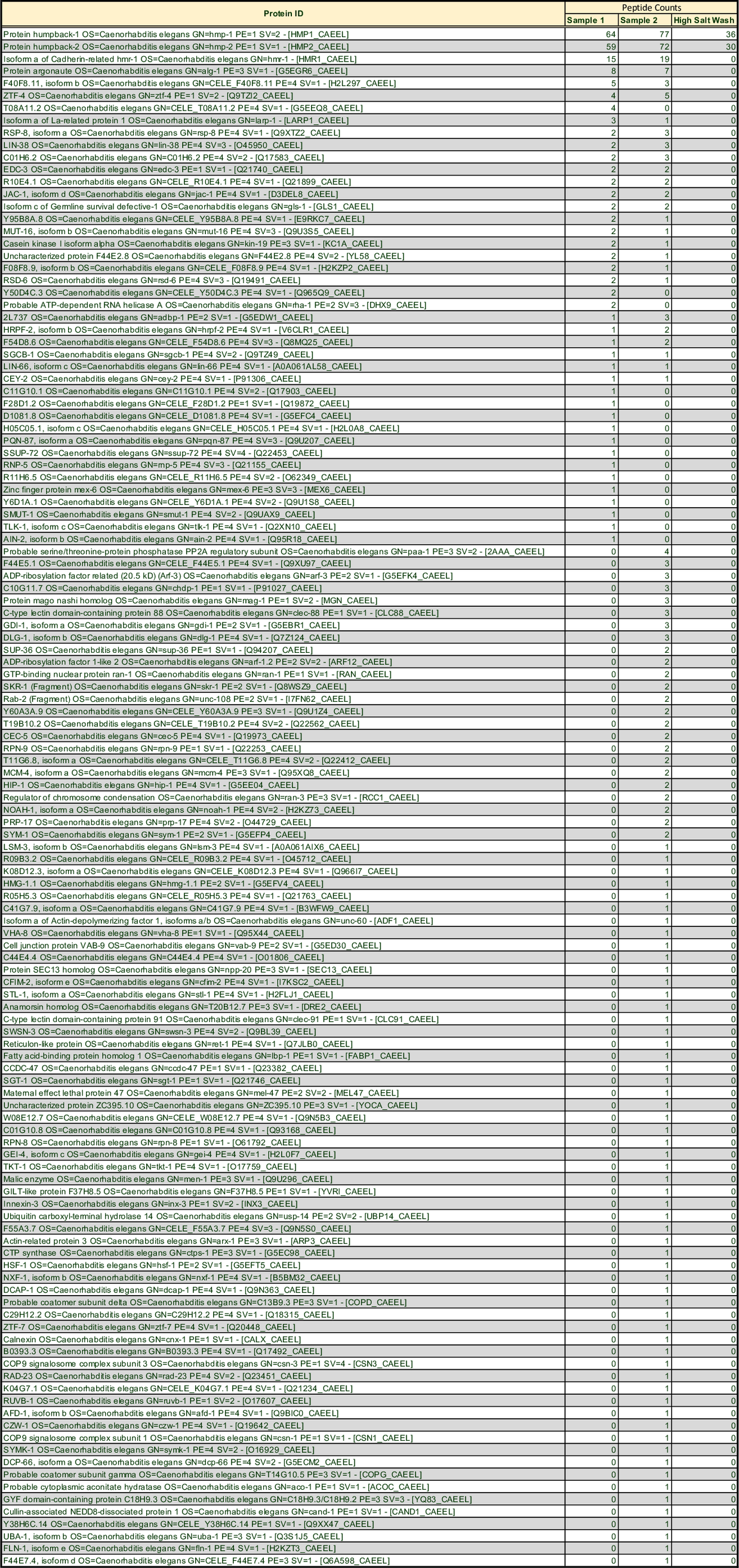
